# A rare mutation in an infant derived HIV-1 envelope glycoprotein alters interprotomer stability and susceptibility to broadly neutralizing antibodies targeting the trimer apex

**DOI:** 10.1101/2020.04.27.065425

**Authors:** Nitesh Mishra, Shaifali Sharma, Ayushman Dobhal, Sanjeev Kumar, Himanshi Chawla, Ravinder Singh, Bimal Kumar Das, Sushil Kumar Kabra, Rakesh Lodha, Kalpana Luthra

**Affiliations:** Department of Biochemistry, All India Institute of Medical Sciences, New Delhi, India; Department of Microbiology, All India Institute of Medical Sciences, New Delhi, India; Department of Pediatrics, All India Institute of Medical Sciences, New Delhi, India

## Abstract

The envelope glycoprotein (Env) of human immunodeficiency virus-1 (HIV-1) is the sole target of broadly neutralizing antibodies (bnAbs). Several mechanisms, such as acquisition of mutations due to the error prone reverse transcriptase, variability of loop length and alterations in glycan pattern are employed by the virus to shield neutralizing epitopes on the env, to sustain survival and infectivity within the host. Identification of mutations that can lead to viral evasion from host immune response is essential for optimization and engineering of Env based trimeric immunogens. Herein, we report a rare leucine to phenylalanine escape mutation (L184F) at the base of hypervariable loop 2 (population frequency of 0.0045%) in a nine-month-old perinatally HIV-1 infected infant broad neutralizer. The L184F mutation disrupted the intramolecular interaction, stabilizing the trimer apex thereby leading to viral escape from autologous plasma bnAbs and known bnAbs, targeting exclusively the N160 glycan at trimer apex and not any other known epitope. The L184F amino acid change led to acquisition of a relatively open trimeric configuration, often associated with tier 1 HIV-1 isolates and an increased susceptibility to neutralization by polyclonal plasma antibodies of weak neutralizers. While there was no impact of the L184F mutation on free virus transmission, a reduction in cell-to-cell transmission was observed. In conclusion, we report a viral escape mutation that plausibly destabilized the trimer apex and favoured evasion from broadly neutralizing antibodies. Such mutations, though rare, should be taken into consideration while designing an immunogen, based on a stable correctly-folded HIV-1 Env trimer.

**Importance:** Design of HIV-1 envelope-based immunogens, capable of eliciting broadly neutralizing antibodies (bnAbs), are currently under active research. Some of the most potent bnAbs target the quaternary epitope at the V2 apex of HIV-1 Env trimer. By studying naturally circulating viruses from an HIV-1 perinatally infected infant, with plasma neutralizing antibodies targeted to the V2-apex, we identified a rare leucine to phenylalanine substitution in two out of six functional viral clones, that destabilized the trimer apex. This single amino acid alteration impaired the interprotomeric interactions that stabilize the trimer apex, resulting in an open trimer conformation, and escape from broadly neutralizing autologous plasma antibodies and known V2-apex directed bnAbs, thereby favouring viral evasion of the early bnAb response of the infected host. Defining the mechanisms by which viral mutations influence the sensitivity of HIV-1 to bnAbs is crucial for the development of effective vaccines against HIV-1 infection.

## Introduction

Elicitation of antibodies capable of neutralizing globally circulating human immunodeficiency virus type 1 (HIV-1) viral variants is one of the vital goals of HIV-1 vaccine research (1, 2). The HIV-1 Envelope glycoprotein (Env) is a trimer of non-covalently linked heterodimers (gp120/gp41)_3_, and is the primary target of broadly neutralizing antibodies (bnAbs). Persistent antigenic stimulation and viral diversification under immune selection pressure are typically associated with the development of bnAbs, though infected infants have been reported to develop bnAbs as early as one-year post-infection (3–5). The bnAbs targeting the viral Env are grouped by epitope class: the variable loop 2 and the N160 glycan (V2-apex), the third variable loop and the N332 glycan (V3/N332-glcan supersite, or the high-mannose patch), the CD4 binding site (CD4bs), gp120/gp41 interface region, the silent face centre, and the membrane proximal external region (MPER) of gp41 (6, 7). These bnAbs are capable of neutralizing diverse circulating variants of HIV-1 and are generated in rare subsets of infected individuals. Passive administration of such bnAbs in animal models and in recently conducted human clinical trials with bnAbs alone, or in combination with antiretroviral therapy, have shown protection from HIV-1 infection (8–13).

No vaccination approach has been successful in inducing bnAbs in humans or standard animal models, though vaccination with native-trimers have thus far induced strain specific and cross-subtype specific nAbs (14–18). To overcome the high level of genetic diversity in HIV-1 Envelope genes, strategies to induce antibodies that cross-react with multiple strains of HIV-1 is the need of the hour. Considerable interest exists in the field relevant to viral features associated with induction and escape mechanisms responsible for V2-apex bnAbs as these bnAbs have been reported to emerge early (19–21), are elicited frequently (22–25), possess relatively low to moderate levels of somatic hypermutations compared to other bnAbs (19–22, 24–26) and show cross-group neutralizing activity with Envs of HIV groups M, N, O, and P (27, 28), thereby identifying the V2-apex as one of the promising Env epitopes for vaccine design. The extraordinary ability of HIV-1 to evade host immunity represents a major obstacle to the development of a protective vaccine. Thus, elucidating the mechanisms employed by HIV-1 to protect its external envelope (Env), which is the sole target of virus-neutralizing antibodies, is an essential step toward developing rational strategies for optimizing Env-based immunogens.

In a recently reported cohort of HIV-1 infected infants with an early plasma bnAb response targeting the Envelope glycoprotein, we had identified a 9-month old infant, AIIMS731, whose plasma bnAbs showed maximum dependence on the V2-apex with 75% breadth at a geometric mean titre (GMT) of 130 against the standardized 12-virus global panel representing global viral diversity (29). In order to understand the virus-antibody dynamics in the context of neutralizing determinants within the V2-apex and early induction of V2-apex targeting plasm bnAbs, herein we studied the viral features associated with escape from plasma bnAbs in an infant broad neutralizer AIIMS731. A rare leucine to phenylalanine mutation (L184F) that impaired the interprotomer stability and consequently led to an open Env trimer conformation was identified. Of note, this rare mutation provided escape from autologous plasma bnAbs as well as several known bnAbs targeting the V2-apex, despite transforming a tier 2 HIV-1 strain into a tier 1 strain, in contrast to known alteration of tiered phenotypes associated with neutralization escape.

## Results

### A rare mutation at the base of hypervariable loop 2 of the viral Env confers resistance to autologous plasma bnAbs in an HIV-1 infected infant broad neutralizer AIIMS731

On the basis of plasma neutralization data against difficult-to-neutralize (Tier 2/3) Global Panel of HIV-1 isolates and epitope mapping done using single base mutants in 25710_2_43, 16055_2_3 and CAP45_G3 and BG505.W6M.C2 pseudoviral backbones (29), an HIV-1 infected infant AIIMS731 was previously categorized as a broad neutralizer with plasma bnAbs targeting the N160-glycan in the V2-apex of HIV-1 Env (***Figure 1A – C***).

**Figure 1.**
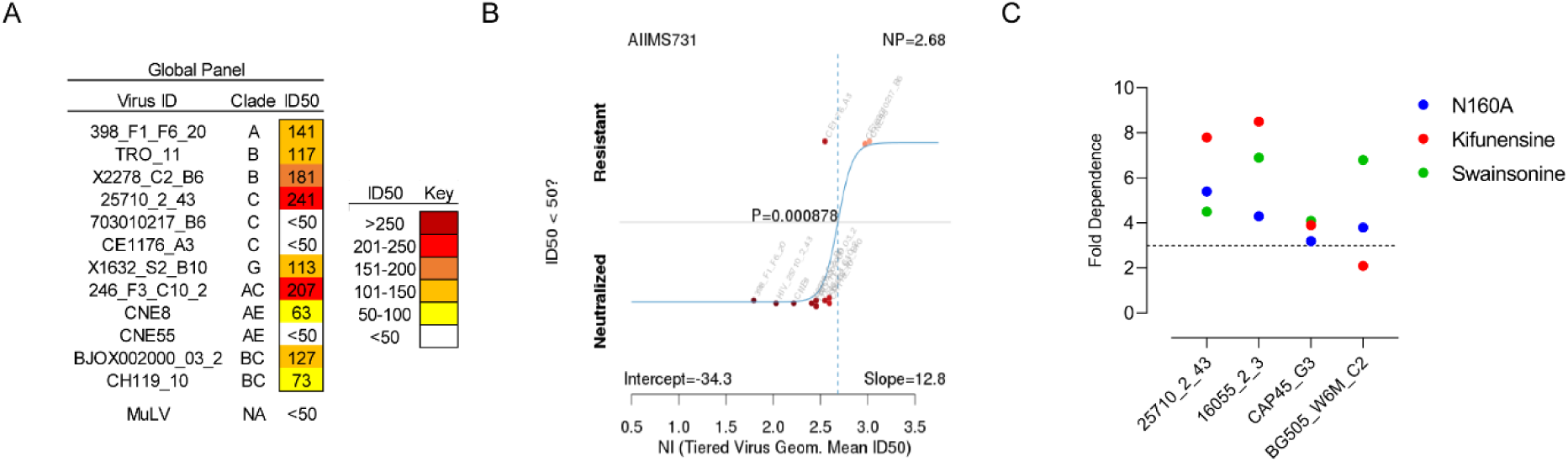
Plasma bnAbs from infant broad neutralizer AIIMS731 target the V2-apex of HIV-1 Env. (A) Heatmap representing HIV-1 specific neutralization titres (inverse plasma dilution) of plasma bnAbs from infant broad neutralizer AIIMS731 against the 12-virus global panel. ID_50_ values are color-coded per the key given, with darker colors implying higher ID_50_ titres. (B) Neutralization indexing explaining the plasma’s ability to neutralize majority of tier 2 viruses on a tier scale that ranks neutralization on a continuous rather than categorical scale. (C) Epitope mapping of AIIMS731 plasma bnAbs showing ID_50_ fold-change against 25710_2_43, 16055_2_3, CAP45_G3 and BG505_W6M_C2 wildtype pseudoviruses and their N160A mutant, as well as pseudoviruses grown in presence of glycosidase inhibitors kifunensine and swainsonine.

In order to evaluate the viral population dynamics associated with the presence of V2-apex plasma bnAbs in this infant broad neutralizer, first, we cloned functional Env genes from the plasma RNA via single genome amplification (SGA), and assessed their susceptibility to autologous plasma bnAbs. A total of 40 Env gene sequences (clade C) from AIIMS731 were available (29), and the depth of SGA sequencing gave us a 90% confidence interval of identifying circulating variants present at a population frequency of 5%. Based on sequence identity, the SGA sequences represented the six dominant R5-tropic circulating strains in AIIMS731 plasma, and were highly homogenous, with sequence variability between the clones ranging from 0.1 to 0.4% (**Figure 2A – C**). Viral variants within the cluster 73105h and 73106f were the dominant circulating strains, while cluster 73105b, 73105c, 73105e and 73105d represented the remaining circulating strains (with population frequency >5%). From each cluster, a single viral variant was cloned into the pcDNA3.1 (+) mammalian expression vector and pseudotyped for neutralization assays.

**Figure 2.**
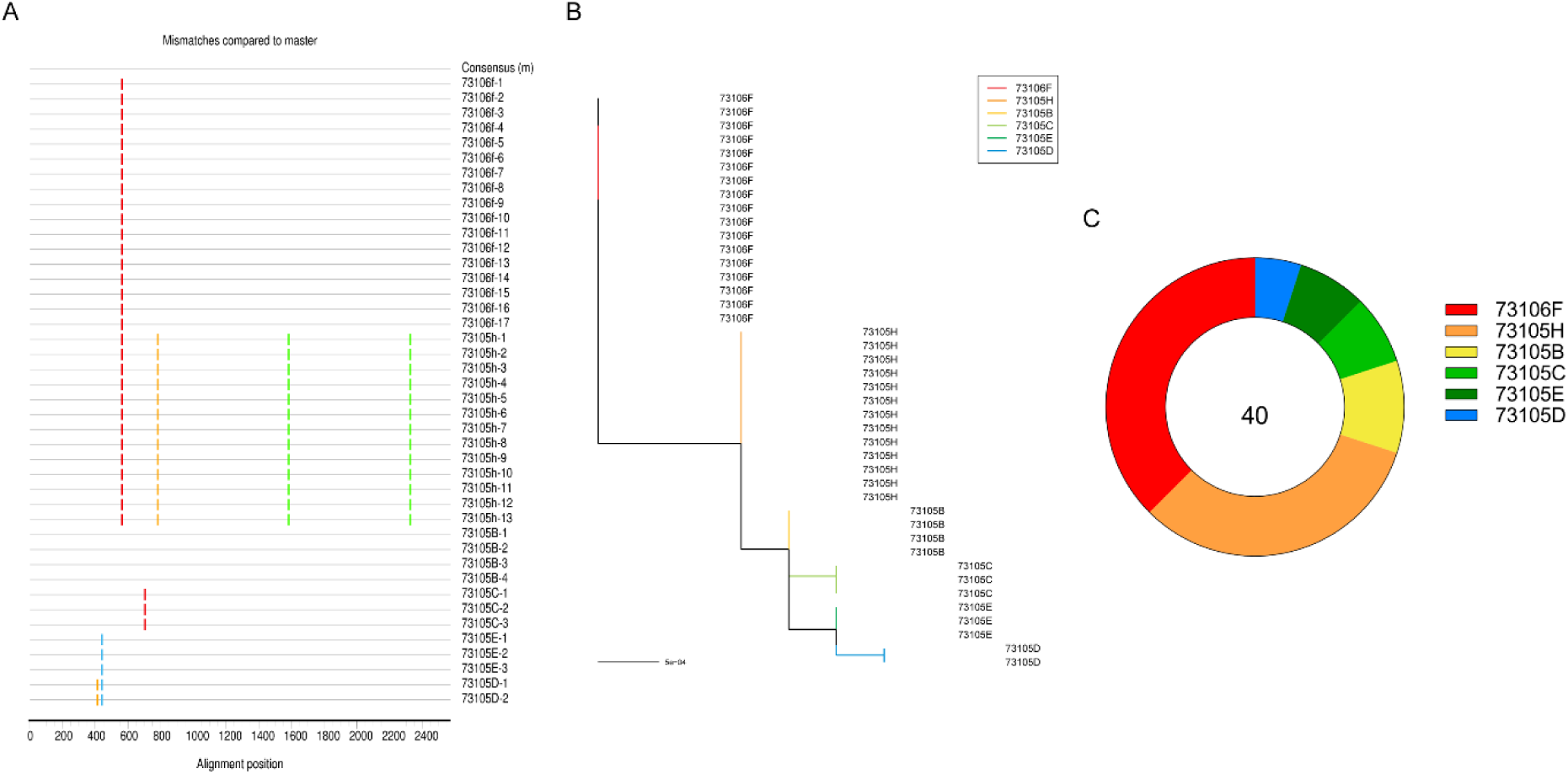
Limited diversity in the circulating viral variants of AIIMS731. (A – B) Highlighter plots with maximum-likelihood trees of 40 SGA env sequences from AIIMS731 showed limited variability in the circulating viral variant, and showed the presence of 6 strains circulating at a population frequency of >5%. In highlighter plot, mutations compared to consensus sequence are represented by green for adenine, blue for cytosine, orange for guanine and red for thymine. In case of 73105h, both of the adenine mutations were silent (Green bars). (C) Donut plot showing the distribution of 40 SGA Env amplicons showed viral variant 73106f and 73105h to be the dominant strains.

Despite the high degree of similarity between the viral variants, a consistent hierarchy of neutralization sensitivity to contemporaneous autologous plasma bnAbs was observed (**Figure 3A**). Viral clone 73105b showed near-complete neutralization (maximum percent neutralization of 87 ± 4%) by autologous plasma bnAbs while clone 73105h and 73106f showed significant abrogation of neutralization sensitivity (maximum percent neutralization of 24 ± 3%) to autologous plasma bnAbs. Despite neutralizing autologous circulating variants with an ID_50_ titres roughly three-fold higher than the median titres against the multiclade panel of HIV-1 isolates (median ID_50_ of 362 vs 127), none of the autologous viruses were completely neutralized by plasma bnAbs with maximum percent neutralization (MPN) for sensitive viruses ranging from 83 to 91%. Both the clones 73105h and 73106f had MPN in the range of 21 to 27%. Of note, plasma bnAb resistant viral variants 73105h and 73106f were the dominant circulating strains (13 and 15 of the 40 SGA sequences, respectively) (**Figure 2C**).

**Figure 3.**
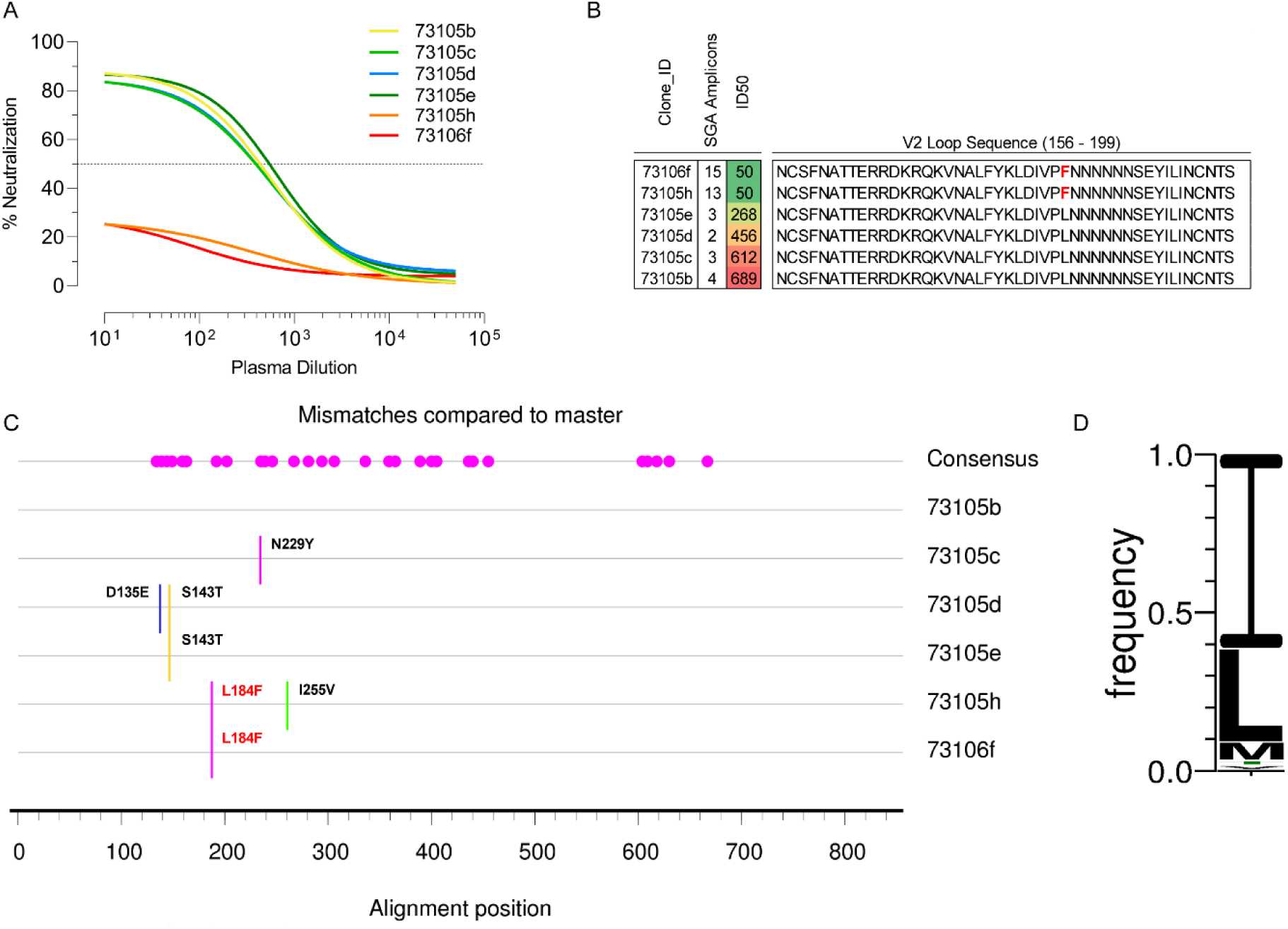
A rare mutation provided neutralization escape from autologous plasma bnAbs in AIIMS731. (A) Neutralization susceptibility of circulating viral variants from AIIMS731 to contemporaneous autologous plasma bnAbs was assessed via neutralization assays based on TZM-bl cells. Even though four of the circulating viral variants were susceptible to plasma bnAbs, maximum percent neutralization ranged from 83 to 91%. Viral variants 73105h and 73106f were resistant to autologous plasma bnAbs (maximum percent neutralization of 24% and 25% respectively). (B) V2-loop sequences (HXB2 numbering 156 – 199) of viral variants showed a leucine-to-phenylalanine mutation at position 184. (C) Complete amino acid sequence comparison between all six viral variants showed a single mutation (L184F) in 73106f (relative to sensitive strain 73105b) that led to neutralization escape from autologous plasma bnAbs. While 73105c also had a rare N229Y mutation, no difference in susceptibility to autologous plasma bnAbs was visible. (D) Amino acid frequency plot at position 184 in all reported HIV-1 Env sequence (7094 sequences) showed abundance of leucine or isoleucine whereas phenylalanine at position 184 occurred at a population frequency of 0.0045% (32/7094 of sequences).

To identify residues conferring resistance to contemporaneous autologous plasma bnAbs, we conducted a comparative sequence analysis. Examination of the core V2-apex bnAb epitope revealed no sequence change despite varying neutralization sensitivity between the viral clones. Of particular note, all the circulating viral variants retained the key epitope-defining patterns of specific amino acid residues and N-linked glycosylation sites in the V2-apex bnAb epitope. Mutations were mapped relative to variant 73105b as it represented the consensus amino acid sequence as well as showed highest susceptibility to autologous plasma bnAbs (**Figure 3B – C**). In case of clone 73105h, in addition to L184F, an additional mutation of I255V (small, non-polar sidechain mutated to another small, non-polar sidechain) within the C2 region was observed. In clone 73105c, N229Y was observed while for clones 73105d (D135E and S143T) and 73105e (S143T), mutations within hypervariable loop 1 (132 – 152, HXB2 numbering) were observed. A single change of L184F was observed in clone 73106f (neutralization resistant) compared to the clone 73105b (most susceptible to neutralization) suggesting that a mutation outside that of the bnAb targeting epitopes in the V2 region may have led to viral escape of the clone 73106f from the V2-apex targeting autologous plasma bnAbs (**Figure 3B – C**). D135E, S143T, and N229Y had no impact on neutralization by autologous plasma bnAb, and were, therefore, excluded from further analysis.

To assess the frequency of the L184F mutation, that led to replacement of a small, non-polar side chain with a bulky non-polar side chain, we examined the variability (amino acid changes). at position 184 (**HXB2 numbering**) in all reported HIV-1 Env sequences Analysis of 7094 Env sequences from Los Alamos National Laboratory (LANL) HIV-1 sequence database revealed the extreme rarity of the L184F mutation. The presence of Phenylalanine at position 184 was found in 32 of the 7094 (a population frequency of 0.0045%) reported viral sequences available at HIV database, with 17 instances reported in clade C (**Table 1**). Position 184 most commonly contains either isoleucine or leucine, at a population frequency of 59.3 and 28.6% respectively (**Table 1 and Figure 3D**). Overall, these results suggested the acquisition of a rare mutation by both 73105h and 73106f viral clone may have led to viral escape from plasma bnAbs in the infected infant AIIMS731.

**Table 1.** Amino acid frequency at position 184. Population frequency for position 184 was calculated using AnalyzeAlign (see methods) for the web alignment of all reported HIV-1 Env sequences (7094) available at LANL HIV database.

| Variant | Count | Frequency |
| --- | --- | --- |
| I | 4206 | 0.5929 |
| L | 2029 | 0.2860 |
| M | 454 | 0.0640 |
| T | 138 | 0.0195 |
| V | 100 | 0.0141 |
| - (Gap) | 50 | 0.0070 |
| F | 32 | 0.0045 |
| S | 20 | 0.0028 |
| N | 16 | 0.0023 |
| D | 11 | 0.0016 |
| A | 9 | 0.0013 |
| P | 6 | 0.0008 |
| E | 4 | 0.0006 |
| R | 4 | 0.0006 |
| * (stop) | 4 | 0.0006 |
| Y | 3 | 0.0004 |
| H | 2 | 0.0003 |
| K | 2 | 0.0003 |
| Q | 2 | 0.0003 |
| C | 1 | 0.0001 |
| G | 1 | 0.0001 |
| W | 0 | 0.0000 |

### L184F provides neutralization escape from V2-apex targeting bnAbs and contributes to the instability of the trimeric form

Next, we evaluated the neutralization susceptibility of all six viral variants from AIIMS731 to assess if the mutations acquired by these viral variants altered neutralization to known V2-apex bnAbs. For all V2-apex bnAbs tested (PG9, PG16, PGT145, PGDM1400, CAP256.25, and CH01), as observed with autologous plasma bnAbs, AIIMS731 viral variants segregated in neutralization sensitive (73105b, 73105c, 73105d and 73105e) and resistant (73105h and 73106f) clusters (**Figure 4**). For resistant variants, we observed a marked increase in IC_50_ and reduction in MPN, with the most significant reduction observed for PG9 and CAP256.25. Except PGDM1400 and CAP256.25, two of the most potent V2-apex bnAbs known, none of the V2-apex bnAbs could reach 100% neutralization, even at higher concentration for 73105b (most sensitive clone). For L184F mutant clones 73105h and 73106f, none of the bnAbs reached 100% neutralization and showed shallow dose-response curves. The slope of neutralization curves for the V2-apex bnAbs were steeper, and had the expected sigmoidal curve for variants in neutralization sensitive cluster compared to L184F mutants 73105h and 73106f (**Figure 4**), suggesting that the viral Envs in sensitive cluster were plausibly homogenously trimeric and uniformly recognized via trimeric-configuration as the V2-apex targeting bnAbs are reported to be trimer-preferring and to target the HIV-1 Env trimer in a closed conformation (30–33).

**Figure 4.**
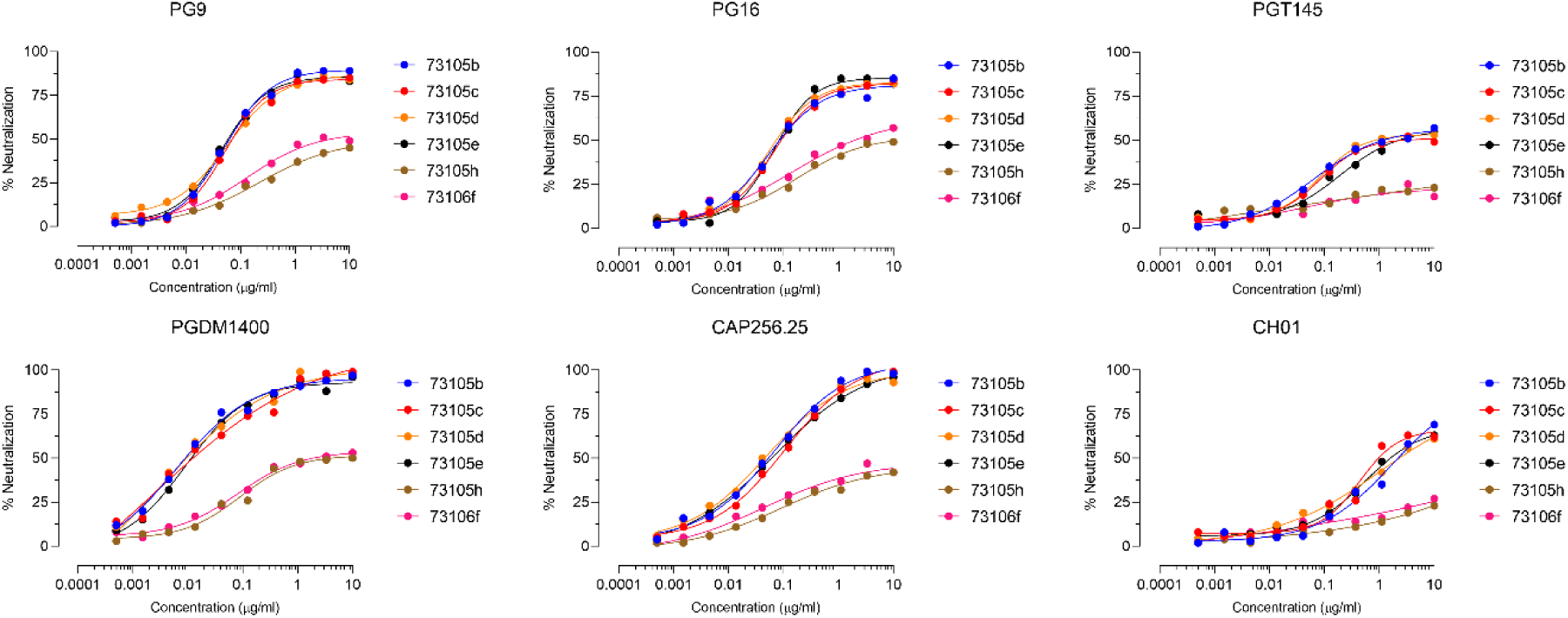
Neutralization curves of AIIMS731 viral variants against V2-apex bnAbs. Neutralization susceptibility of all six viral variants to the V2-apex bnAbs (PG9, PG16, PGT145, PGDM1400, CAP256.25 and CH01) was assessed via neutralization assays based on TZM-bl cells. Of note, except for PGDM1400 and CAP256.25, none of the V2-apex bnAbs reached 100% neutralization for AIIMS731 autologous plasma bnAbs sensitive viral cluster (73105b, 73105c, 73105d and 73105e) while with the L184F mutant clones 73105h and 73106f (autologous plasma bnAb resistant cluster), all V2-apex bnAbs showed markedly lower neutralization efficiency. For 73106f, maximum neutralization of 57% and 53% was reached with PG16 and CAP256.25 respectively. Neutralization assays were repeated thrice, and curves were drawn based on average neutralization.

Next, we used an exhaustive panel of bnAbs targeting other known epitopes on the Env in order to assess the neutralization efficiency of bnAbs other than those targeting the V2 region, by comparing the neutralization curves for all six viral variants. The bnAb panel consisted of: V3/N332-glycan supersite bnAbs (10-1074, BG18, AIIMS-P01, PGT121, PGT128 and PGT135); CD4bs bnAbs (VRC01, VRC03, VRC07-523LS, N6, 3BNC117, and NIH45-46 G54W); silent face targeting bnAb (PG05); fusion peptide and gp120/gp41 interface bnAbs (PGT151, 35O22 and N123-VRC34.01); and MPER bnAbs (10E8, 4E10 and 2F5). Neutralization assays showed similar neutralization phenotypes of these bnAbs for all six variants, regardless of their susceptibility to V2-apex bnAbs (**Table 2**), suggesting that the L184F mutation was specific to the viral escape from susceptibility to neutralization by V2-apex bnAbs and had negligible effect on the neutralization by other classes of bnAbs.

**Table 2.**
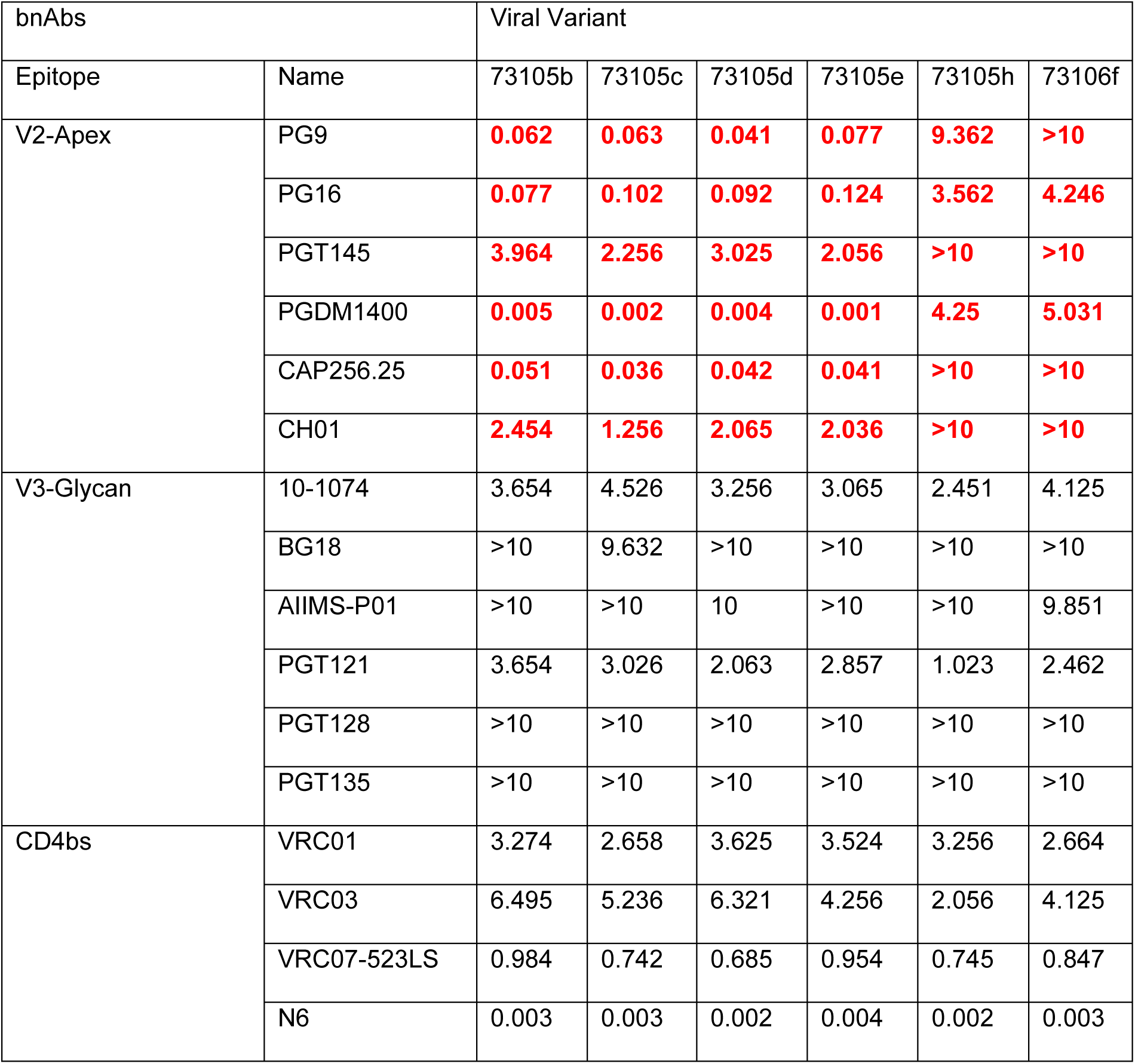
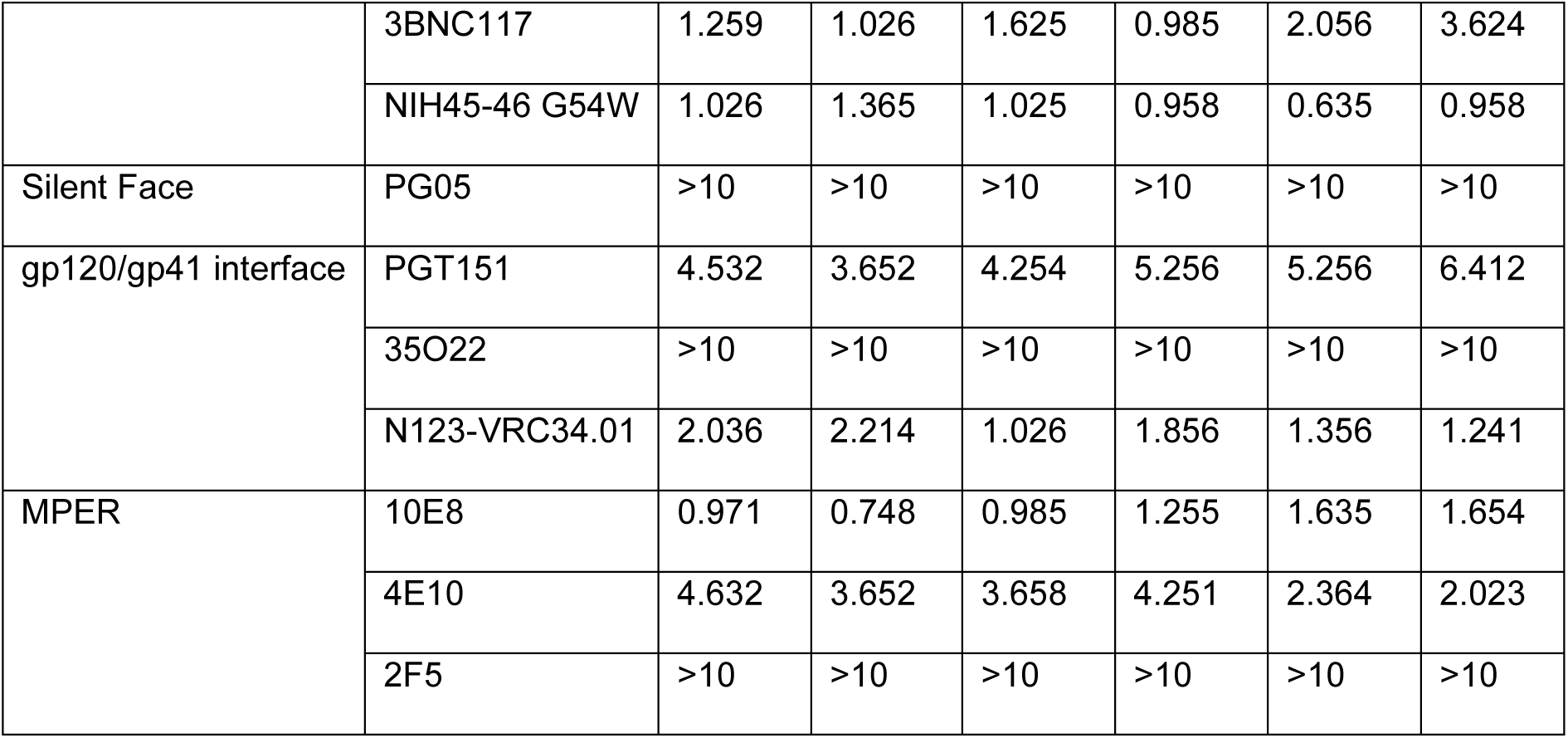
Neutralization of 73105b and 73106f by known bnAbs. Neutralization susceptibility of 73105b and 73106f were assessed utilizing a broad panel of bnAbs targeting all major antigenic sites on HIV-1 Env. IC50 values (50% inhibitory concentration) and MPN (maximum percent neutralization) for all tested bnAbs are shown and grouped according to the antigenic sites (V2-Apex, V3/N332-glycan supersite, CD4bs, Silent Face, gp120/gp41 interface, MPER). Neutralization assays were performed with TZM-bl cells and repeated thrice. IC50 values and MPN were calculated based on average neutralization.

| bnAbs |  | Viral Variant |  |  |  |  |  |
| --- | --- | --- | --- | --- | --- | --- | --- |
| Epitope | Name | 73105b | 73105c | 73105d | 73105e | 73105h | 73106f |
| V2-Apex | PG9 | 0.062 | 0.063 | 0.041 | 0.077 | 9.362 | >10 |
|  | PG16 | 0.077 | 0.102 | 0.092 | 0.124 | 3.562 | 4.246 |
|  | PGT145 | 3.964 | 2.256 | 3.025 | 2.056 | >10 | >10 |
|  | PGDM1400 | 0.005 | 0.002 | 0.004 | 0.001 | 4.25 | 5.031 |
|  | CAP256.25 | 0.051 | 0.036 | 0.042 | 0.041 | >10 | >10 |
|  | CH01 | 2.454 | 1.256 | 2.065 | 2.036 | >10 | >10 |
| V3-Glycan | 10-1074 | 3.654 | 4.526 | 3.256 | 3.065 | 2.451 | 4.125 |
|  | BG18 | >10 | 9.632 | >10 | >10 | >10 | >10 |
|  | AIIMS-P01 | >10 | >10 | 10 | >10 | >10 | 9.851 |
|  | PGT121 | 3.654 | 3.026 | 2.063 | 2.857 | 1.023 | 2.462 |
|  | PGT128 | >10 | >10 | >10 | >10 | >10 | >10 |
|  | PGT135 | >10 | >10 | >10 | >10 | >10 | >10 |
| CD4bs | VRC01 | 3.274 | 2.658 | 3.625 | 3.524 | 3.256 | 2.664 |
|  | VRC03 | 6.495 | 5.236 | 6.321 | 4.256 | 2.056 | 4.125 |
|  | VRC07-523LS | 0.984 | 0.742 | 0.685 | 0.954 | 0.745 | 0.847 |
|  | N6 | 0.003 | 0.003 | 0.002 | 0.004 | 0.002 | 0.003 |
|  | 3BNC117 | 1.259 | 1.026 | 1.625 | 0.985 | 2.056 | 3.624 |
|  | NIH45-46 G54W | 1.026 | 1.365 | 1.025 | 0.958 | 0.635 | 0.958 |
| Silent Face | PG05 | >10 | >10 | >10 | >10 | >10 | >10 |
| gp120/gp41 interface | PGT151 | 4.532 | 3.652 | 4.254 | 5.256 | 5.256 | 6.412 |
|  | 35O22 | >10 | >10 | >10 | >10 | >10 | >10 |
|  | N123-VRC34.01 | 2.036 | 2.214 | 1.026 | 1.856 | 1.356 | 1.241 |
| MPER | 10E8 | 0.971 | 0.748 | 0.985 | 1.255 | 1.635 | 1.654 |
|  | 4E10 | 4.632 | 3.652 | 3.658 | 4.251 | 2.364 | 2.023 |
|  | 2F5 | >10 | >10 | >10 | >10 | >10 | >10 |

In neutralization assays performed with non-neutralizing antibodies (non-nAbs) targeting the V3 loop (447-52d and 19b) and CD4-induced epitopes (17b, A32, 48d, b6), L184F mutants 73105h and 73106f showed weak neutralization by the V3 loop non-nAbs (MPN of 29% and 28% for 447-52D; 29% and 31% for 19b, respectively) and CD4i non-nAbs (MPN of 35% and 36% for 17b; 26% and 23% for 48d respectively), although even at high concentrations, none of the non-nAbs reached IC_50_ titres against the L184F mutant (**Figure 5A – B**). Of note, 17b binds preferentially to the CD4-induced, CCR5 co-receptor binding site epitope on Env (34). Thus, L184F mutation resulted in increased susceptibility to neutralization by antibodies known to target the relatively more open conformation of the Env, suggesting that the rare L184F mutation allowed the Env to sample more open states (characteristics of Tier 1A and 1B viral variants) resembling CD4-bound conformation where the CCR5 binding site is exposed; though this observation can only be accurately validated by undertaking in-depth structural studies.

**Figure 5.**
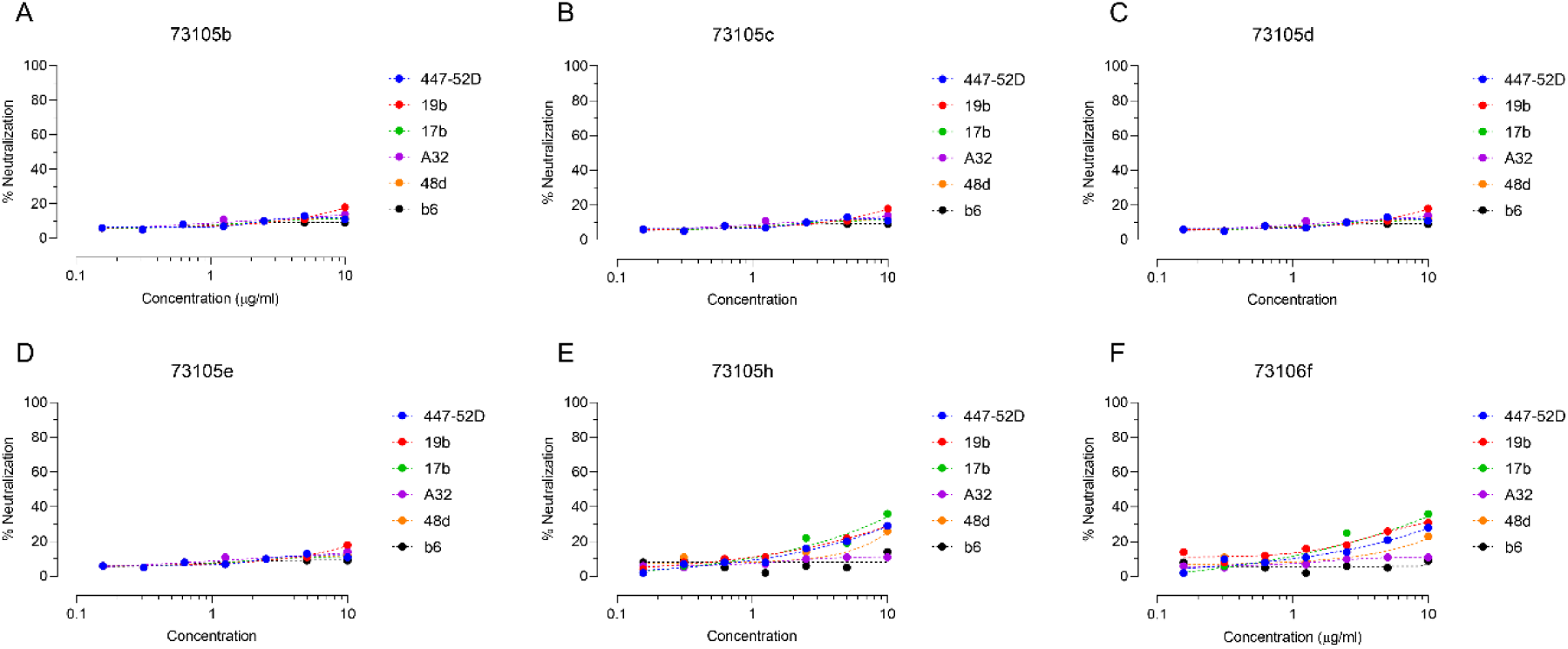
Neutralization of AIIMS731 viral variants by non-nAbs targeting the V3 Loop and CD4-induced epitopes. (A – F) Neutralization susceptibility of all six viral variants to the V3 loop targeting non-nAbs (447-52D and 19b) and CD4-induced non-nAbs (17b, A32, 48d and b6) was assessed via neutralization assays based on TZM-bl cells. Viral variant 73105h and 73106f showed moderate neutralization by 447-52D, 19b, 17b and 48d. Neutralization assays were repeated thrice, curves were drawn, and MPN were calculated based on average neutralization.

Taken together, these results suggest L184F mutation conferred resistance to neutralization via trimer-preferring V2-apex bnAbs, and allowed the Env trimer to transition towards a more open configuration that partially exposed the occluded non-nAbs epitopes within V3 loop and CD4bs.

### Preferential recognition of the closed Env trimer by potent plasma antibodies from pediatric neutralizers

Destabilization of the trimer apex has been shown to alter the neutralization susceptibility of HIV-1 Env to antibodies present in the plasma of infected individuals. As L184F mutation resulted in a more open trimer configuration, we next evaluated the sensitivity of L184F mutant to a panel of HIV-1 clade C infected pediatric patient plasma with varied neutralization potency (weak versus strong neutralizers) against the global panel of representative HIV-1 isolates (35–37). Patient plasma antibodies neutralizing more than half the global panel were considered strong neutralizers while those neutralizing less than half the panel were considered weak to moderate neutralizers depending on their breadth and potency. The neutralization susceptibility profile of L184F mutants 73105h and 73106f to plasma antibodies of well-characterized HIV-1 clade C infected pediatric donors (whose plasma antibodies showed varied neutralization activity against the 12-virus global panel) showed comparable ID_50_ values between weak and strong neutralizers (**Figure 6A – C**), confirming that viral variant belonging to autologous plasma bnAb neutralization sensitive cluster (73105b, 73105c, 73105d and 73105e) showed a Tier 2 phenotype while the viral variants 73105h and 73106f had a Tier 1B neutralization phenotype.

**Figure 6.**
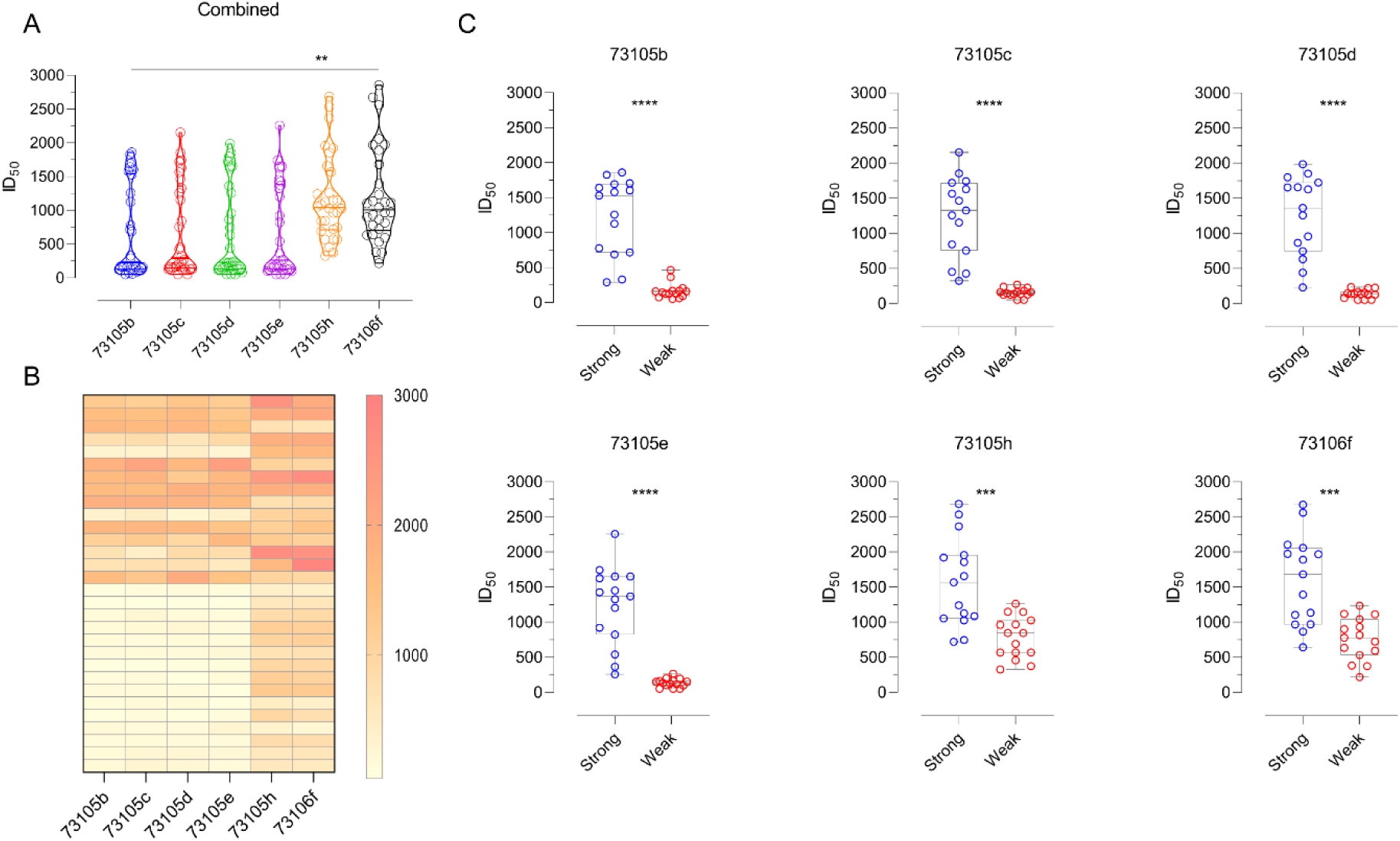
Viral variant 73106f is highly susceptible to subtype-matched heterologous plasma antibodies. (A – B) Violin plot and heatmap representing the neutralization susceptibility of AIIMS731 circulating viral variants against plasma antibodies from HIV-1 infected pediatric individuals in chronic stages of disease was assessed. Distinct neutralization profile was seen for AIIMS731 autologous plasma bnAbs sensitive (73105b, 73105c, 73105d and 73105e) and resistant (73105h and 73106f) viral clusters. The plasma panel contained well-characterized HIV-1 clade C infected pediatric donors whose plasma antibodies showed varied neutralization activity against the 12-virus global panel. Plasmas were categorized as strong or weak based on their breadth and potency against the 12-virus global panel (see methods). Comparison is shown for 73105b and 73106f, though similar patterns were observed on comparing sensitive vs. resistant clusters. (C) Viruses belonging to sensitive cluster were primarily neutralized by plasma samples that were categorized as strong while viruses belonging to resistant cluster showed considerable neutralization by plasma samples categorized as weak. P-values are given by asterisks where ** implies p-value <0.01, *** implies p-value <0.001 and **** implies p-value <0.0001.

The 73105b (most sensitive to autologous plasma bnAbs) virus clone was highly susceptible to the plasma of strong neutralizers with ID_50_ values ranging from 1:248 to 1:1965 while the susceptibility to weak neutralizers ranged from 1:50 to 1:188 (1:50 was the lowermost dilution tested). For the L184F mutant 73106f, ID_50_ titres of strong neutralizers (range – 1:388 to 1:2756) versus weak neutralizers (range – 1:176 to 1:1246) had similar profile suggesting regardless of their ability to generate antibodies capable of targeting closed Env trimers, weak neutralizers develop high titres of antibodies targeting the open configuration of the Env trimer.

### L184F escape mutation resulted in reduced entry kinetics in cell-cell transmission

As V1V2 stabilizes the Env spike forming the trimer apex, we next performed a stability-of-function assay, called T90 assay, that determines Env stabilization (viral infectivity) as a function of temperature (38, 39), to evaluate the effect of the L184F mutation on the Env stability. A slight increase in T90 value, the temperature at which the viral infectivity decreased by 90% in 1 hour, from 43.27 to 43.69 (p = 0.19) was observed, though the impact of L184F on thermal stability did not appear markedly noticeable (**data not shown**).

Changes at the trimer apex have been shown to alter virus sensitivity and often come with a fitness cost (39–42). In order to investigate the effect of acquisition of the rare L184F viral immunotype on the functional stability of the Env, we assessed the impact of L184F mutation on viral infectivity in free virus and cell-cell transmission. The relative infectivity of all six viral variants was assessed by titration curves after normalizing pseudoviral infectivity by using the viral stock dilution that gave a relative luminescence unit (RLU) of 150,000 in TZM-bl cells. No change in the infectivity of the L184F mutant was observed in case of infection with free virus, though we detected substantial variability in entry kinetics of L184F mutants 73105h and 73106f in cell-cell transmission (**Figure 7A – B**), suggesting a plausible fitness cost associated with escape from plasma bnAbs via the acquisition of rare viral L184F immunotype, that needs to be further confirmed.

**Figure 7.**
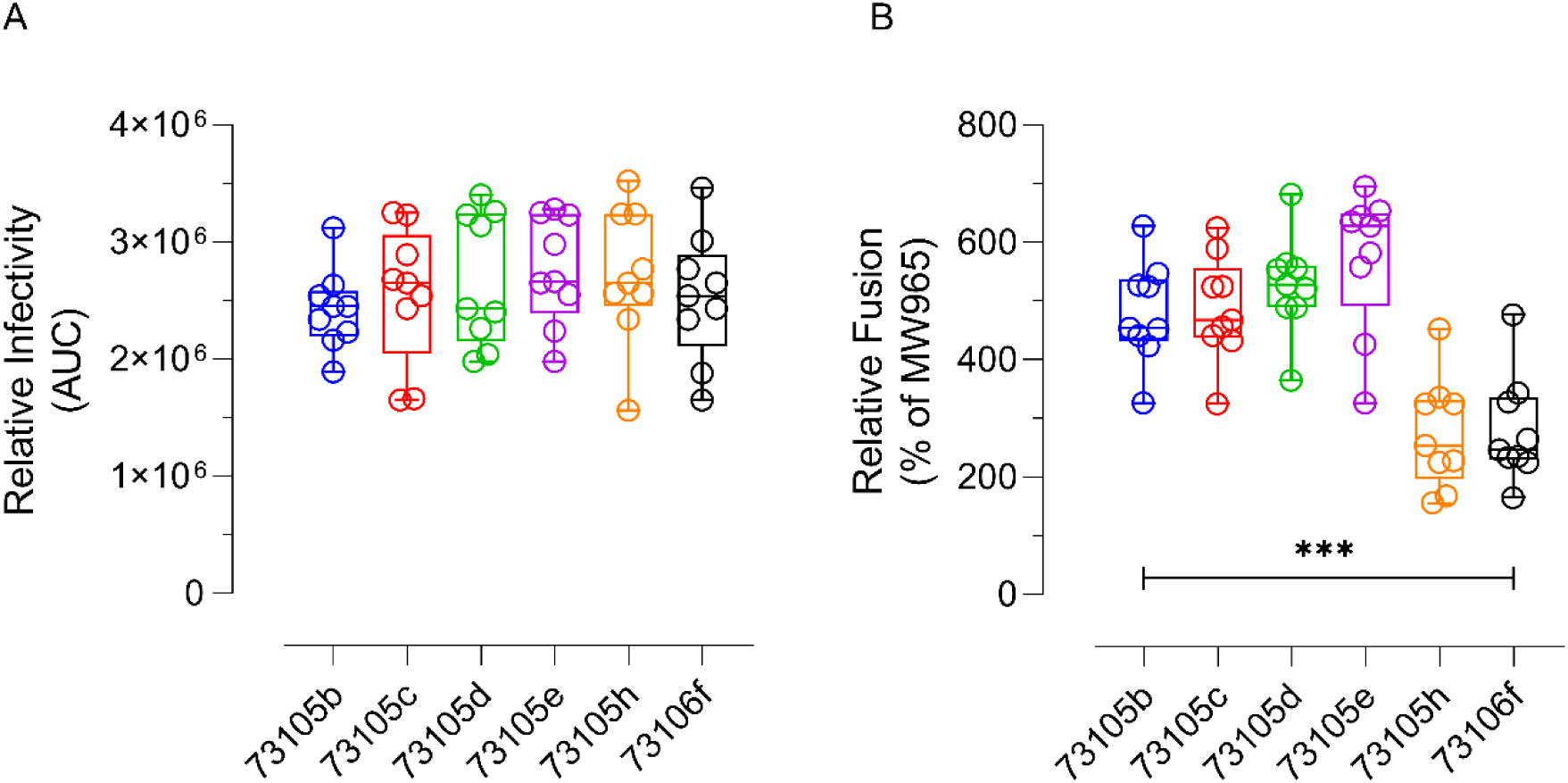
L184F mutation led to reduced cell-cell transmission. (A) Pseudoviruses were titrated after normalization and replicate titration curves were used to calculate the area under the curve (AUC) values. Each experiment was repeated thrice in triplicates providing a total of 9 reference values. (B) Fusogenicity in co-cultures of Tat/Env co-transfected 293T and TZM-bl cells were used as a measure of cell-to-cell transmission ability. Fusion of AIIMS731 Env in relation to fusion observed with well-characterized Env of HIV-1 isolate MW965.26 was calculated. Each experiment was repeated thrice in triplicates providing a total of 9 reference values. 2-tailed student’s t test was used for comparison (*** implies p-value <0.001). Comparison is shown for 73105b and 73106f, though similar patterns were observed on comparing sensitive vs. resistant clusters.

### L184F escape mutation impaired interprotomer interaction at trimer apex

The V2-apex bnAbs target quaternary epitopes formed by interprotomeric interactions at the apex of HIV-1 Env trimer. The core epitope for V2-apex bnAbs is formed by the N-linked glycan sites N156 and N160, and the lysine rich region of strand C (HXB2 numbering: 156 – 177). As the L184F escape mutation did not arise within the core epitope, and that this mutant virus showed a tier 1 neutralization phenotype, we reasoned that the L184F mutation was plausibly responsible for disrupting the interprotomer interactions that stabilize the close conformation of the Env trimer.

To elucidate the mechanism by which the L184F mutation could have disrupted the conformation of the Env trimer, we analyzed the L184F mutation using the ligand-free pre-fusion closed structure of BG505 SOSIP.664 HIV-1 Env trimer (PDB ID: **4ZMJ**). Residues 165 and 184 in BG505 were changed to their counterpart (R165 and L184) in 73105b. In previous reports, I184 of one protomer has been shown to interact with L165 of the neighbouring protomer. This inter-protomeric interaction has been shown to be critical for quaternary interactions leading to the stabilization of the V1V2 regions of neighbouring protomers, and its loss has been shown to render JR-FL, a clade B HIV-1 strain, highly sensitive to V3 mAbs (42, 43). On similar lines, we observed L184 of one promoter interacting with R165 of another protomer [via van der Waals interactions between the solvent-accessible surface (SAS) of R165 and L184 on neighbouring protomers] (**Figure 8A**).The side chain of L184 was observed to be outward facing and did not make significant intra-protomeric interactions. On mutating L184 to F184, disruption of SAS between the bulky side chain of F184 on one protomer and R165 on neighbouring protomer was seen (**Figure 8B**). In addition, we generated a homology model based on the sequence of 73105b based on multiple structural templates, and after loop refinement, Man-9-Glycan sites were added to potential-N-linked glycosylation sites (PNGS) in silico to produce a near-fully glycosylated gp160 trimeric model. In 73105b homology model, similar interprotomeric interactions were seen between L184 of one protomer with R165 of neighbouring protomer which were lost when L184 was mutated to F184.

**Figure 8.**
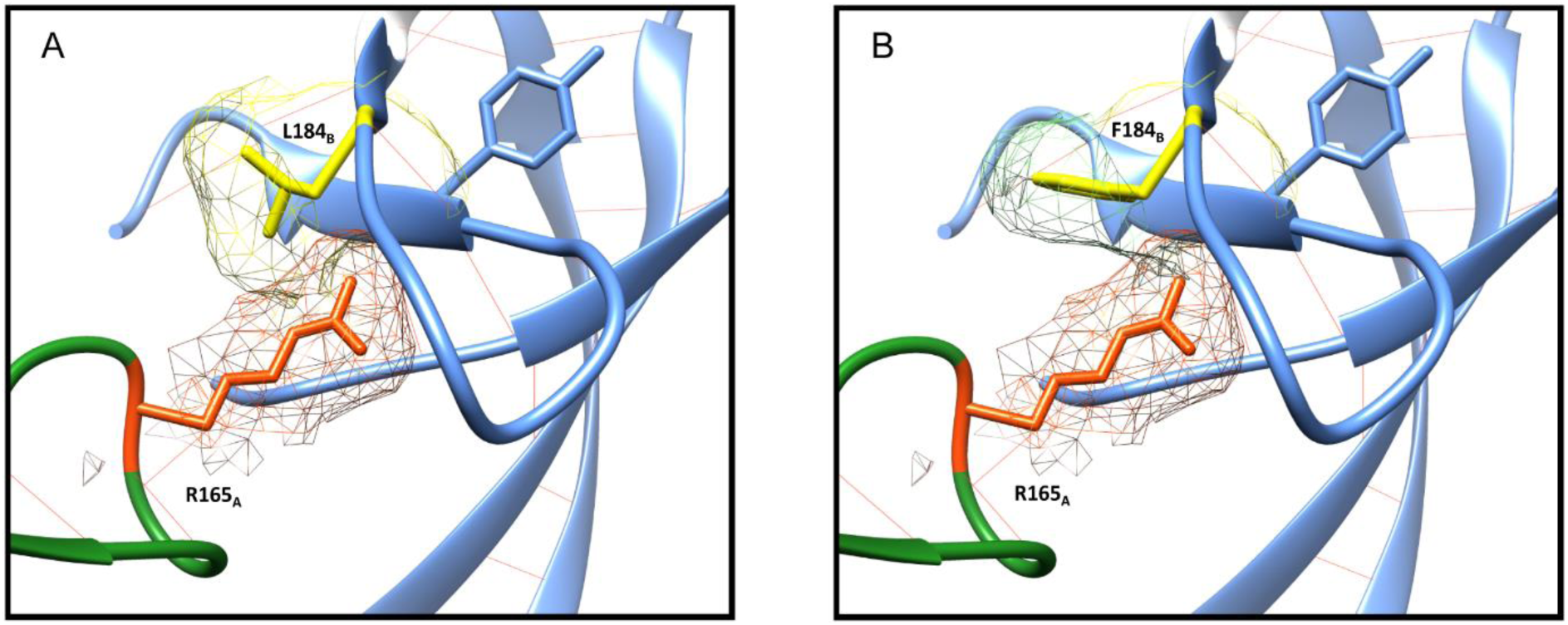
Critical role of L184 in modulating interprotomer interactions at the trimer apex. Interprotomer interactions between R165 (protomer A) and L184 (protomer B). The dot mesh surrounding R165_A_ (Orange) and L184_B_ and F184_B_(Yellow) represent the solvent-accessible surfaces (SAS) (van der Waals surfaces expended by the water molecule radius). In (A) Interprotomer contacts (lipophilic) between R165_A_ and L184_B_ can be seen by overlapping SAS. In case of F184_B_, no interprotomer contacts can be seen, evident by the lack of SAS overlap. The HIV-1 Env protomeric backbones are represented by two distinct colors (protomer A as green and protomer B as cornflower blue). The illustration was generated from PDB entry 4ZMJ. L165 and I184 were rotamerized to respective residues in 73105b (R165 and L184) and 73106f (R165 and F184).

## Discussion

During HIV-1 infection, the humoral immune response targets the HIV Envelope (Env) glycoprotein which consists of three heavily glycosylated non-covalently linked gp41-gp120 protomers (3–7). While strain-specific antibodies recognize exposed and variable sites, bnAbs target relatively conserved and occluded sites, including the quaternary V1V2 epitope at the trimer apex (V2-Apex), V3/N332-glycan supersite, CD4-binding site (CD4bs), gp120-gp41 interface, and membrane proximal external region (MPER). Of these, bnAbs targeting the quaternary V1V2 epitope (called V2-apex bnAbs) are elicited frequently and relatively early (19–26). HIV-1 eludes recognition by host bnAbs through a variety of mechanisms, though the most common mechanism includes sequence alterations that can lead to large variations in sensitivity to antibody-mediated neutralization among different circulating viral isolates. Herein, we investigated the viral escape mechanisms in a 9-month-old perinatally HIV-1 infected infant with broadly neutralizing plasma antibodies targeting the V2-apex.

Escape from contemporaneous autologous plasma bnAbs occurred by a rare leucine to phenylalanine mutation at position 184. Interestingly, a high degree of similarity was observed between circulating viral strains, regardless of their sensitivity to plasma bnAbs, and L184F mutation, alone, was enough for escape from neutralization by plasma bnAbs. A similar neutralization profile for sensitive and resistant strains were observed when susceptibility to reported bnAbs targeting diverse epitopes on the Env was assessed. While the L184F mutation did not alter neutralization profile of the viral strains to bnAbs targeting the V3/N332 glycan supersite, CD4-binding site, gp120/gp41 interface or MPER, a significant reduction in neutralization susceptibility to several V2-apex bnAbs was seen. The most significant reduction observed was for CAP256.25 (also referred to as VRC25.26), a trimer-specific bnAb that recognizes HIV-1 env trimers via its long protruding loop that interacts with strand C, insertions into the apex hole at trimer 3-fold axis as well as electrostatic interactions with cationic V1V2 surface residues (19, 30, 33). Given the complex mode of trimer recognition via CAP256.25 which is a combination of PG9 and PGT145 class of bnAbs, and therefore, its stringent need for a closed conformation of Env trimer (33), the L184F mutation most likely seemed to alter the conformation of the Env trimer.

As V2-apex bnAbs target the closed conformation of the trimer, based on neutralization profile observed, we hypothesized that the most plausible reason for the loss of susceptibility to V2-apex bnAbs was the loss of closed conformation of the Env. In its closed conformation, the Env trimer occludes several immunodominant epitopes that are targeted by non-neutralizing antibodies (non-nAbs) (39, 41). When neutralization profile against several non-nAbs (those targeting CD4-induced epitopes and V3 loop) was assessed, the L184F mutant virus showed relatively higher neutralization susceptibility to non-nAbs. Of note, none of the non-nAbs could achieve 50% neutralization (17b, which binds preferentially to the CD4-induced, CCR5 co-receptor binding site epitope on Env achieved a maximum percent neutralization of 32%) (34), suggesting that the L184F mutation did not substantially alter the conformation of the Env. In addition, the L184F mutant virus showed high susceptibility to neutralization by polyclonal antibodies present in the plasma of weak neutralizers (HIV-1 infected patients that do not generate a potent response to HIV-1 Env) (35–37). HIV-1 Envs are categorized into tiers based on neutralization susceptibility to plasma antibodies (44, 45). Conventionally, viruses categorized as Tier 1 (generally lab-adapted strains) are easier to neutralize by plasma antibodies, while Tier 2 viruses are substantially more resistant. Tier 3 strains exhibit exceptional resistance to antibody-mediated neutralization. The neutralization tier phenotypes of HIV-1 isolates can be understood in the context of the dynamic nature of Env trimers on the virus surface (34, 46). These trimers spontaneously transition between closed, open, and at least one intermediate conformation. Open trimers expose more epitopes than closed trimers, and are typically reported to have a Tier 1 neutralization phenotype while trimers in closed state have Tier 2/3 neutralization phenotype. Tier 3 Envs have been shown to exclusively exist in a relatively narrow range of closed conformation. Overall, our findings suggest that the L184F led to the acquisition of a relatively open trimeric configuration, which is most typically associated with Tier 1 HIV-1 isolates. HIV-1 Env evades recognition by antibodies through diverse mechanisms, but one of the most effective mechanism is the adoption of a closed trimeric configuration (typically associated with Tier 2 or 3 Envs). Our observation of acquisition of a rare mutation that led to an open, yet inaccessible to plasma V2-apex bnAbs, Env state suggests HIV-1 can lose its trimeric Env form to evade plasma bnAbs.

In silico structural analysis was then utilized to reveal the molecular feature that conferred resistance to V2-apex bnAbs. Substitution of the small side chain of leucine with the bulky non-polar side chain of phenylalanine led to disruption of interprotomer interactions that have been observed to be critical for maintenance of the trimeric apex. Mutations that alter or disrupt interprotomer contacts in Env trimer have been shown to change its susceptibility to several classes of bnAbs (39, 41–43). Overall, our results provide information on the role of a rare escape mutation L184F in the viral envelope that led to resistance to bnAbs targeting the V2-apex. Understanding the impact of viral escape mutations on the sensitivity of HIV-1 to bnAbs provides vital information for optimization of vaccines candidates against HIV-1. Furthermore, our data is suggestive of the prominent role of L184 mediated intramolecular interactions that are necessary for the maintenance of the trimer apex and adds to the information on conformational epitopes on the HIV-1 envelope towards the development of effective vaccines.

## Materials and methods

### Study design and participants

The current study was designed to assess the viral population dynamics in an infant broad neutralizer with plasma bnAbs targeting the V2-apex of HIV-1 Env. At the time of recruitment, AIIMS731 was antiretroviral naïve and asymptomatic; was 9-months old with a CD4 count of 1385 cells/mm^3-^ and viral load on log scale was 5.894 RNA copies/ml (785,000 RNA copies/ml); was recruited from the Pediatric Chest Clinic, Department of Pediatrics, AIIMS. After written informed consent from guardians, blood was drawn in 3-ml EDTA vials, plasma was aliquoted for plasma neutralization assays, viral RNA isolation, and viral loads. The study was approved by institute ethics committee of All India Institute of Medical Sciences (IECPG-307/07.09.2017).

### Plasmids, viruses, monoclonal antibodies, and cells

Plasmids encoding HIV-1 env genes representing different clades, monoclonal antibodies and TZM-bl cells were procured from NIH AIDS Reagent Program. 10-1074 and BG18 expression plasmids were kindly provided by Dr. Michel Nussenzweig, Rockefeller University, USA; VRC07-523LS and N123.VRC34.01 expression plasmids were provided by Dr. John Mascola, VRC, NIH, USA; and PG05 expression plasmids were provided by Dr. Peter Kwong, VRC, NIH, USA. CAP256.09, CAP256.25 and b6 were procured from IAVI Neutralizing Antibody Centre, USA. 293T cells were purchased from the American Type Culture Collection (ATCC).

### HIV-1 envelope sequences and phylogenetic analysis

HIV-1 envelope genes were PCR amplified from plasma viral RNA by single genome amplification and directly sequenced commercially. Individual sequence fragments of SGA amplified amplicons were assembled using Sequencher 5.4 (Gene Code Corporation). Subtyping for SGA sequences was performed with REGA HIV subtyping tool (400bp sliding window with 200bp steps size). Inter-clade recombination was examined with RIP 3.0 (Recombinant Identification Program) and with jpHMM. Nucleotide sequences were aligned with MUSCLE in MEGA X. Maximum-likelihood trees were computed with MEGA X using a general-time reversal substitution model incorporating a discrete gamma distribution with 5 invariant sites.

### Cloning of autologous HIV-1 envelope genes and production of replication incompetent pseudoviruses

Autologous replication incompetent envelope pseudoviruses were generated from AIIMS731 as described previously (29, 37). Briefly, viral RNA was isolated from 140 µl of plasma using QIAamp Viral RNA Mini Kit, reverse transcribed, using gene specific primer OFM19 (5’ - GCACTCAAGGCAAGCTTTATTGAGGCTTA – 3’) and Superscript III reverse transcriptase, into cDNA which was used in two-round nested PCR for amplification of envelope gene using High Fidelity Phusion DNA Polymerase (New England Biolabs). The envelope amplicons were purified, and ligated into pcDNA3.1D_ITR vector via overlap extension cloning. Pseudoviruses were prepared by co-transfecting 1.25 µg of HIV-1 envelope containing plasmid with 2.5 µg of an envelope deficient HIV-1 backbone (PSG3Δenv) vector at a molar ratio of 1:2 using PEI-MAX as transfection reagent in HEK293T cells seeded in a 6-well culture plates. Culture supernatants containing pseudoviruses were harvested 48 hours post-transfection, filtered through 0.4µ filter, aliquoted and stored at −80°C until further use. TCID_50_ was determined by infecting TZM-bl cells with serially diluted pseudoviruses in presence of DEAE-Dextran, and lysing the cells 48 hours post-infection. Infectivity titres were determined by measuring luminescence activity in presence of Bright Glow reagent (Promega).

### Infectivity and neutralization assay

Viral infectivity and neutralization assays were carried out using TZM-bl cells, a genetically engineered HeLa cell line that constitutively expresses CD4, CCR5 and CXCR4, and contains luciferase and β-galactosidase gene under HIV-1 tat promoter, as described before (37). Viral infectivity was determined after normalizing pseudoviruses to an RLU value of 150,000 followed by titration curve generation to calculate relative infectivity. Neutralization studies included heat-inactivated plasmas from AIIMS731 and previously characterized thirty plasmas of chronically infected children (35–37), 25 bnAbs (PG9, PG16, PGT145, PGDM1400, CAP256.25, CH01, 10-1074, BG18, AIIMS-P01, PGT121, PGT128, PGT135, VRC01, VRC03, VRC07-523LS, N6, 3BNC117, NIH45-46 G54W, PG05, PGT151, 35O22, N123.VRC34.01, 10E8, 4E10 and 2F5) and 6 non-nAbs (447-52D, 19b, 17b, A32, 48d, b6). Briefly, envelope pseudoviruses were incubated in presence of serially diluted heat inactivated plasmas, bnAbs or non-nAbs for one hour. After incubation, freshly Trypsinized TZM-bl cells were added, with 25 µg/ml DEAE-Dextran. The plates were incubated for 48h at 37°C, cells were lysed in presence of Bright Glow reagent, and luminescence was measured. Using the luminescence of serially diluted bnAbs or plasma, a non-linear regression curve was generated and titres were calculated as the bnAb concentration, or reciprocal dilution of serum that showed 50% reduction in luminescence compared to untreated virus control. For epitope mapping, 25710_2_43, 16055_2_3, CAP45_G3 and BG505_W6M_C2 N160K mutants and pseudoviruses grown in presence of kifunensine and swainsonine were used and greater than 3-fold reduction in ID_50_ titres were classifies as dependence.

### HIV-1 Env stability-of-function assay

Thermostability (T90) assays were performed as described previously (38, 39). Briefly, 73105b and 73106f pseudotyped viruses were incubated at temperatures from 37°C to 50°C for 60 minutes using temperature gradient on PCR thermal cycler (BioRad). Pseudoviruses were then aliquoted in a 96-well culture plate, followed by addition of 10,000 TZM-bl cells per well. Infectivity was determined by performing titration curves and plotted as a function of temperature. T90 values were interpolated as the temperature at which virus infectivity decreased by 90%.

### HIV-1 Env cell-to-cell fusion assay

HIV-1 Env mediated cell-to-cell fusion assays were performed as described previously (47, 48). Briefly, 293T cells were co-transfected with pTAT (pcTat.BL43.CC, #11785, NIH AIDS Reagent Program) and envelope plasmid encoding for either 73105b, 73106f and MW965.26 Env. To remove intra and inter-assay variability, all technical replicates were co-transfected and expressed on the same day. 293T cells transfected with pTAT only were used as negative control. 24-hour post-transfection, 10,000 pTAT/Env or pTAT transfected 293T cells were mixed with 10,000 TZM-bl cells (1:1 ratio) in 96-well culture plates, and incubated for 6 hours. Luciferase activity was measured using Bright Glow reagent and fusion activity (cell-to-cell transmission) for 73105b and 73106f was normalized to MW965.26 Env mediated fusion.

### Structural modelling and analysis

Crystal structure of ligand-free BG505.SOSIP.664 HIV-1 Env trimer (PDB ID: **4ZMJ**) was used to assess the interprotomeric interactions due to L184 and F184. Mutations were modelled using the Rotamers tool with Dunbrack 2010 rotamer library (49). For modelled trimers based upon AIIMS731 Env sequences, crystal structure of HIV-1 envelope trimer (PDB ID: **6P65, 6PWU, 6MZJ** and **6B0N**) were used as template to generate homology models based on 73105b amino acid sequence. Homology modelling was carried out using the Modellar 9.22 interface (50) with UCSF Chimera package (51). High-mannose (Man-9) glycans were added to the modelled trimer using the glycoprotein builder interface available at Glycam web server (http://glycam.org/). To limit the computational complexity, Man-9 glycans were selected as computational complexity increases exponentially with complex and/or oligomannose glycans which have multiple branching topologies.

### Statistical analysis

2-tailed student’s t test for paired analysis and Mann-Whitney U test for unpaired analysis were used. For assessing the relative infectivity, area under curves were calculated. All statistical analyses were performed on GraphPad Prism 8. A p-value of <0.05 was considered significant.

### Data and materials availability

The SGA amplified HIV-1 envelope sequences used for inference of phylogeny and highlighter plots are available at GenBank with accession numbers MT366192 – MT366197. All data required to state the conclusions in the paper are present in the paper. Additional information related to the paper, if required, can be requested from the authors.

## Acknowledgments

We thank study subject AIIMS731 for participating in this study. We are thankful to NIH AIDS Reagent program for providing HIV-1 envelope pseudovirus plasmids, bnAbs, non-nAbs and their expression plasmids, and TZM-bl cells, and Neutralizing Antibody Consortium (NAC), IAVI, USA for providing bnAbs. We are thankful to Dr. Michel Nussenzweig for providing 10-1074 and BG18 bnAb expression plasmids; Dr. John Mascola for providing VRC07-523LS and N123.VRC34.01 bnAb expression plasmids; and Dr. Peter Kwong for providing PG05 bnAb expression plasmids.

## Author contributions

N.M designed the study, performed SGA amplification, pseudovirus cloning, and neutralization assays, analyzed data, wrote the initial manuscript, revised and finalized the manuscript. S.S and A.D contributed to SGA amplification, pseudovirus cloning, and neutralization assays. S.K and H.C expressed PGDM1400, CAP256.25, BG18, 10-1074 and AIIMS-P01 bnAb. R.S, B.K.D, R.L, and S.K.K, provided the samples from HIV-1 infected infant AIIMS731. R.L and S.K.K provided patient care and management. S.S, A.D, S.K, and H.C edited and revised the manuscript. K.L designed the study, edited, revised and finalized the manuscript.

